# Flowtrace: simple visualization of coherent structures in biological fluid flows

**DOI:** 10.1101/086140

**Authors:** William Gilpin, Vivek N. Prakash, Manu Prakash

## Abstract

We present a simple, intuitive algorithm for visualizing time-varying flow fields that can reveal complex flow structures with minimal user intervention. We apply this technique to a variety of biological systems, including the swimming currents of invertebrates and the collective motion of swarms of insects. We compare our results to more experimentally-diffcult and mathematically-sophisticated techniques for identifying patterns in fluid flows, and suggest that our tool represents an essential “middle ground” allowing experimentalists to easily determine whether a system exhibits interesting flow patterns and coherent structures without the need to resort to more intensive techniques. In addition to being informative, the visualizations generated by our tool are often striking and elegant, illustrating coherent structures directly from videos without the need for computational overlays. Our tool is available as fully-documented open-source code available for MATLAB, Python, or ImageJ at www.flowtrace.org.

## 2 Introduction

Flow visualization is an essential technique for studying biological processes arising in many diverse areas, including ecology,^1^ biomechanics,^2^ and molecular biology.^3, 4^ Observations of the trajectories of many tracer particles—whether fluorescent beads, microbubbles, vesicles inside a moving cell, or flocking bacteria—can reveal rich information about the way that biological motion such as molecular transport, ciliary flows, or active force generation is coordinated over many length and timescales.^5–8^ Additionally, flow visualization aids in characterization and secondary validation of many standard techniques, such as microfluidics and flow cytometry^9, 10^

The simplest qualitative flow visualization techniques involve observations of the motion of passive scalars, like dyes or smoke, as they are advected by the flow. These experiments have the benefit of being relatively straightforward to perform, and can often yield immediate insight into the global structure and mixing properties of a flow.^11^ However, at length and time scales dominated by diffusive effects, or in flows characterized by large separations in the timescales of different processes, the results of such studies can be difficult to interpret.^12, 13^ As a result, in many contexts quantitative flow characterization techniques based on the motion of tracer particles are preferable.^14–16^

However, standard quantitative flow visualization techniques—particle tracking and particle image velocimetry (PIV)—are suffciently diffcult to implement and optimize that rigorous fluid dynamical visualization techniques remain prohibitive in many experimental contexts.^13^ While PIV and related techniques have been widely-applied and optimized for certain systems, such as the study of fish swimming^17^ or bloodflow mechanics,^8^ in less-established contexts these techniques require specialized modifications of apparatuses such as laser light sheets or point probes to be constructed.^4, 18^ As a result, system-dependent techniques are often necessary,^19^ particularly when particle motion is only partially visible in the data, such as from image streaks^9^ or out of focus drift due to limited depth of field.^20^ This issue is even more prominent for flows in non-traditional media, such as in the collective motion of flocks and herds.^21–23^

Nonetheless, many standard concepts in fluid mechanics—such as vortices, jets, and turbulence—are widely-known to researchers throughout the sciences, who may recognize the likely presence of these features in their data even without the need for quantitative flow characterization tools.^24, 25^ This suggests that additional flow visualization tools are necessary for systems for which dye-based qualitative techniques are unavailable, but quantitative tracer-particle studies are unnecessary.

Here, we present Flowtrace, an intuitive, qualitative visualization technique that can reveal the presence of various flow structures in experimental videos, allowing it to be used either as a primary analysis tool for presence/absence studies or to motivate the use of more complicated flow quantification techniques. Our technique is based purely on image processing of the input data, rather than numerical reconstruction of scalar fields like vorticity, allowing it to be used as a straightforward “first pass” characterization technique for biological systems where traditional techniques are either unnecessary to support qualitative observations, or prohibitively difficult due to the length and timescales involved. While our technique is straightforward and intuitive, we were unable to find previous reports of it in the literature—and, importantly, we find that it has surprising utility for studying patterns in a wide range of biological data sets.

## 3 Method

### 3.1 Algorithm

Our algorithm generalizes a common technique for generating long-exposure photographs from videos, in which the maximum (or minimum) intensity projection of a time series is taken in order to generate pathlines for bright objects moving against a dark background— resulting in “motion streaks” across the image.^5^ This technique has previously been used to create star trails, a popular astronomical visualization generated by taking the maximum intensity projection of a stabilized video of the night sky.^26^

Our primary extension of this technique is to take a single long video, and then take the maximum intensity projection of small groups of successive frames, in order to generate sequential images showing pathlines at different times. The resulting time series of pathline images shows how the shapes of the pathlines change over time. Concretely, for a 100 frame video, a Flowtrace video with 30-frame traces consists of the maximum intensity projection of frames 1-31, 2-32, 3-33, etc, and so forth. The sequence ends when frames 70-100 are projected, resulting in a 70 frame video. In addition to this basic operation, various other operations can be combined with the maximum intensity projection operator in order to yield improved results. The process is illustrated graphically in Figure 1C.

Symbolically, let each frame of the movie be a vector of pixel values and locations, *v*_*ij*_ [*t*], where *t* = 1, 2, …, *N* represents the index of a frame in a video consisting of *N* frames, and *i* and *j* denote the coordinates of a pixel in the image. Suppose that the tracer particles are brightly-colored objects moving against a dark background. In this case, we define a series of maximum-intensity projections, **p**[*t*], in terms of a forward convolution operator,

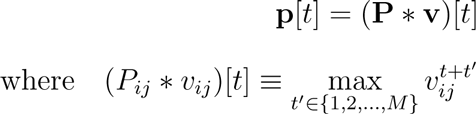

where *M* < *N* is some subset of the frames in the video. As the time index *t* “slides” forward across successive indices 1, 2, 3, …, the maximum intensity projection is taken across successive runs of *M* frames that each differ by two images (the first and the last). This results in a set of maximum intensity projections, 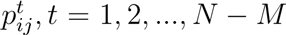, that constitute a new video generated from the original data set. Importantly, the number of frames in the generated video (*N* − *M*) is almost equal to the number of frames in the original video (*N*). In this paper, we refer to each subsequence of *M* images as a *substack*, and the sequence of positions taken by a single particle moving across *M* frames as a *pathline*. The parameter *M*, the timescale over which particle pathlines are visualized in each frame, represents the only parameter that the user must specify in order to use the tool.

During convolution, the projection operator **P** “slides” across the entire sequence of frames, operating on the video in overlapping sets of *M* frames. This operator can be composed with other pre-processing operations in order to achieve different effects; in the code described below, other operations defined include median subtraction (to remove slowly-moving objects), color inversion (for dark objects moving against a lighter background), differential weighing (coloring or darkening each frame in the *M* frame sequence a different amount, in order to show a gradient across the pathlines indicating time), and pairwise differencing (to isolate objects that move faster than 1 px/frame). Each of these operations symbolically represents composition before convolution, such that the final image series is ((**P** º **G**) * **v**)[*t*], where **G** is the pre-processing operation.

### 3.2 Software package and options

Our algorithm is implemented as “Flowtrace,” an open-source package for MATLAB, Python, or ImageJ at www.flowtrace.org. Full tutorials and sample image sets are provided there. Table 1 summarizes the primary user-specified arguments and options available for the code; optional arguments are passed as a *struct* object in MATLAB, as keyword arguments in Python, and as checkboxes in a GUI for ImageJ.

Oftentimes a dataset features two well-separated velocity scales; one for tracer particles and another slow component for gradual drift, bulk flow, etc. In such cases Flowtrace performs best when the projection length *M* is set so that the fast particles travel far within the field of view, while objects moving at the slower speed move relatively little. This is true for the pathlines shown for the two feeding current-generating organisms shown in Figure 1, which are drawn for projection periods that are long compared to the transit time of tracer particles, but short compared to the gradual motion of each organism’s body. As a results the features of each organism’s anatomy remain sharp in the image.

In some cases, these timescales are not well separated (resulting in motion blur for slowly-moving objects), or there are stationary objects and obstacles in the image that obscure the pathlines. For these situations, it is useful to apply a background subtraction operation to each substack before performing the projection. For objects moving slowly relative to the tracer particles, but fast enough to exhibit noticeable motion blur, the most aggressive background option is “take_difference”, which takes the pairwise differences among all consecutive images before applying the projection. However, if some tracer particles move slowly relative to others, these will also vanish from the image. For nearly-stationary background objects, “subtract_first” or “substract_median” yield similar results based on the types of objects.

Occasionally it is convenient to highlight the directionality of time in the resulting pathlines, particularly when still frames from the output time series are used for analysis. In this case, directionality can be indicated by applying a color gradient across time using the optional argument “color_series”, or by applying a linear intensity gradient using the optional argument “fade_tails”.

### 4 Results and Discussion

We have applied Flowtrace to a variety of biological systems, and it proves surprisingly effective in illustrating the complicated dynamics of unsteady flows arising in a variety of contexts. By varying the projection time interval, *τ* (and thus the pathline lengths), we have studied biological phenomena occurring over a wide range of length and time scales. Methods for each of the experiments briefly described below are discussed further in the Supplementary Materials.

**Table 1.** Options and parameters for Flowtrace. Full documentation for individual versions of Flowtrace for ImageJ, Python, and MATLAB can be found at www.flowtrace.org

| Required Argument (type) | Description |
| --- | --- |
| frames_to_merge (integer) | The number of input frames to combine per output frame |
| image_dir (string) | The location to save output files |
| Optional Argument (type) | Description (default) |
| invert_color (boolean) | For images comprising dark objects moving against a light background (false) |
| subtract_median (boolean) | Subtract the median of the substack from the substack before projection (false) |
| take_difference (boolean) | Take the pairwise differences of images before projection (false) |
| subtract_first (boolean) | Subtract the first image of each substack from the substack before projection (false) |
| add_first (boolean) | Add the first image of each substack to each projected image (false) |
| color_series (boolean) | Apply a color gradient across pathlines (false) |
| frames_to_skip (integer) | Number of alternate frames to omit from each projection (0) |
| use_parallel (boolean; Python only) | Use multithreading across cores (false) |
| max_cores (integer; Python only) | The maximum number of threads to use when running in parallel (4) |
| fade_tails (boolean; MATLAB only) | Apply an intensity gradient across pathlines (false) |

Figure 1A shows three representative frames from a Flowtrace movie of a starfish larva, which in previous work we have shown creates dynamic vortex arrays around its body as it continuously adjusts its feeding currents.^27^ The full Flowtrace movie from which the video was generated shows the diffierent vortex patterns smoothly transitioning into one another over time, indicating how these different, dynamic flow patterns evolve due to the animal’s control mechanisms (see Supplementary Video 11 of the referenced paper).^27^ In Figure 1B we explore similar ciliary flows generated during filter feeding by the protozoan *Stentor sp*. (size ~ 50*µm*). The videos and images (*τ* = 3 s) capture the helical motion of algae particles as the organism slowly rotates its stalk (Supplementary Video S1).

In the sea anemone *Aiptasia pallida* (~ 1 mm), an inverted-color video (*τ* = 4 min) shows the breakup of a water jet as the animal peristaltically pumps water containing tracer beads into its body cavity (Figure 2C, Supplementary Video S3). In a similar video of a moon snail veliger (~ 1 mm), Flowtrace processes a color video generated by a DSLR camera by projecting each channel separately, generating a true color video of the formation of the dipolar flow field created by the swimming animal (Figure 2D, Supplementary Video S4).

In addition to passive fluid tracer particles, Flowtrace can be applied to active particles and ecological data. In a crawling army ant swarm (~ 5 mm, filmed in the field with a handheld iPhone), Flowtrace shows the convergence of the ants’ paths (*τ* = 1 s, Figure 2B, Supplementary Video S2). Similarly, in a ~ 1 m wide swarm of flying midges, applying the optional parameter color series shows color gradients (representing time) along the midges’ pathlines, allowing the tightening of the flock to be visualized (Figure 2A, Supplementary Video S1, raw data taken from the referenced paper).^28^

The simple sliding projection technique used by Flowtrace appears to be largely-unknown in the biological sciences and fluid dynamics literature, despite the ease with which it can be implemented. Flowtrace can reproduce the core qualitative conclusions of several studies, including our recent study on larval starfish swimming,^27,29^ in which the key observation of distinct feeding and swimming vortex arrays generated by the animals is first determined based on Flowtrace videos. We have found that our method works particularly well for studies of feeding currents because there is a wide separation between the timescales of advection and field variation, such that tracer particles have sufficient time to map out the structure of the flow field before it undergoes further variation. Moreover, feeding phenomena typically involve small length scales and long timescales, for which traditional dye advection visualization techniques would fail due to rapid diffusive mixing.

**Figure 1.**
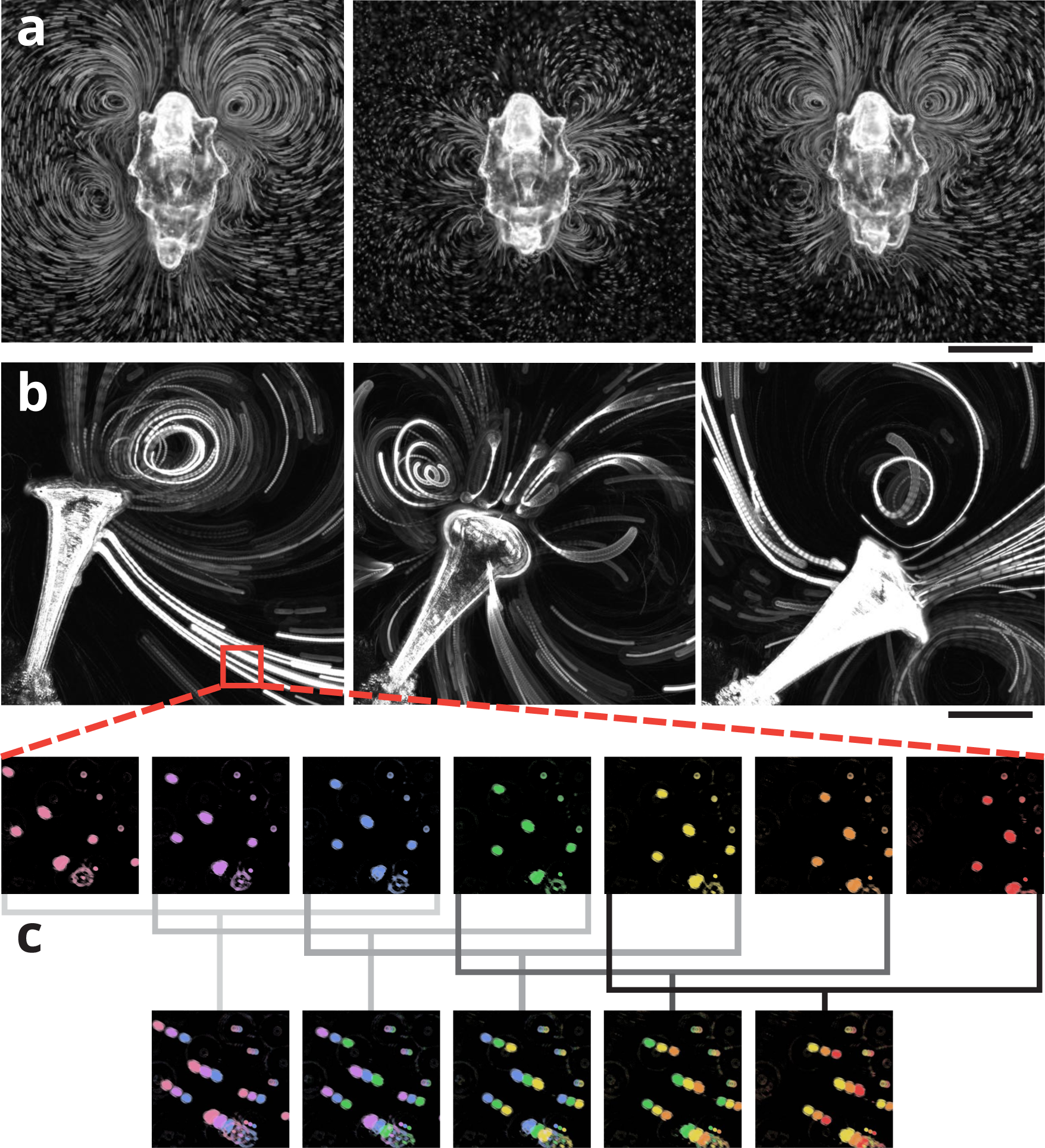
The Flowtrace algorithm. (**a.**) Three stills from a video of feeding currents generated by the larva of the starfish *Patiria miniata*. The full video is available as Supplementary Video 11 of our previous work.^27^ (**b.**) The gyration of *Stentor sp.* as it filters water containing 6 *µ*m beads (*τ* = 3 s, timepoints = 0, 6.5, 18 s, scale bar 175 *µ*m). (**c.**) A false-color detail from panel b illustrating the “sliding projection” used by Flowtrace to generate pathlines (Supplementary Video 1; *τ* = 3 frames, frame rate = 20 fps, scale bar 25 *µ*m).

We have compared Flowtrace to other techniques for identifying structures in fluid flows, and we find that Flowtrace can qualitatively reproduce the results of more-sophisticated dynamical analysis using either vorticity contours or finite-time Lyapunov exponents^30, 31^ (see Supplementary Analysis). Our tool is available in effcient, multithreaded implementations for Fiji/ImageJ, Python 2 and 3, and MATLAB. We envision a broad community—from microscopists to ecologists to fluid physicists—will find Flowtrace useful, and so the full source code and documentation is available at http://www.flowtrace.org or for pull requests on GitHub.

**Figure 2.**
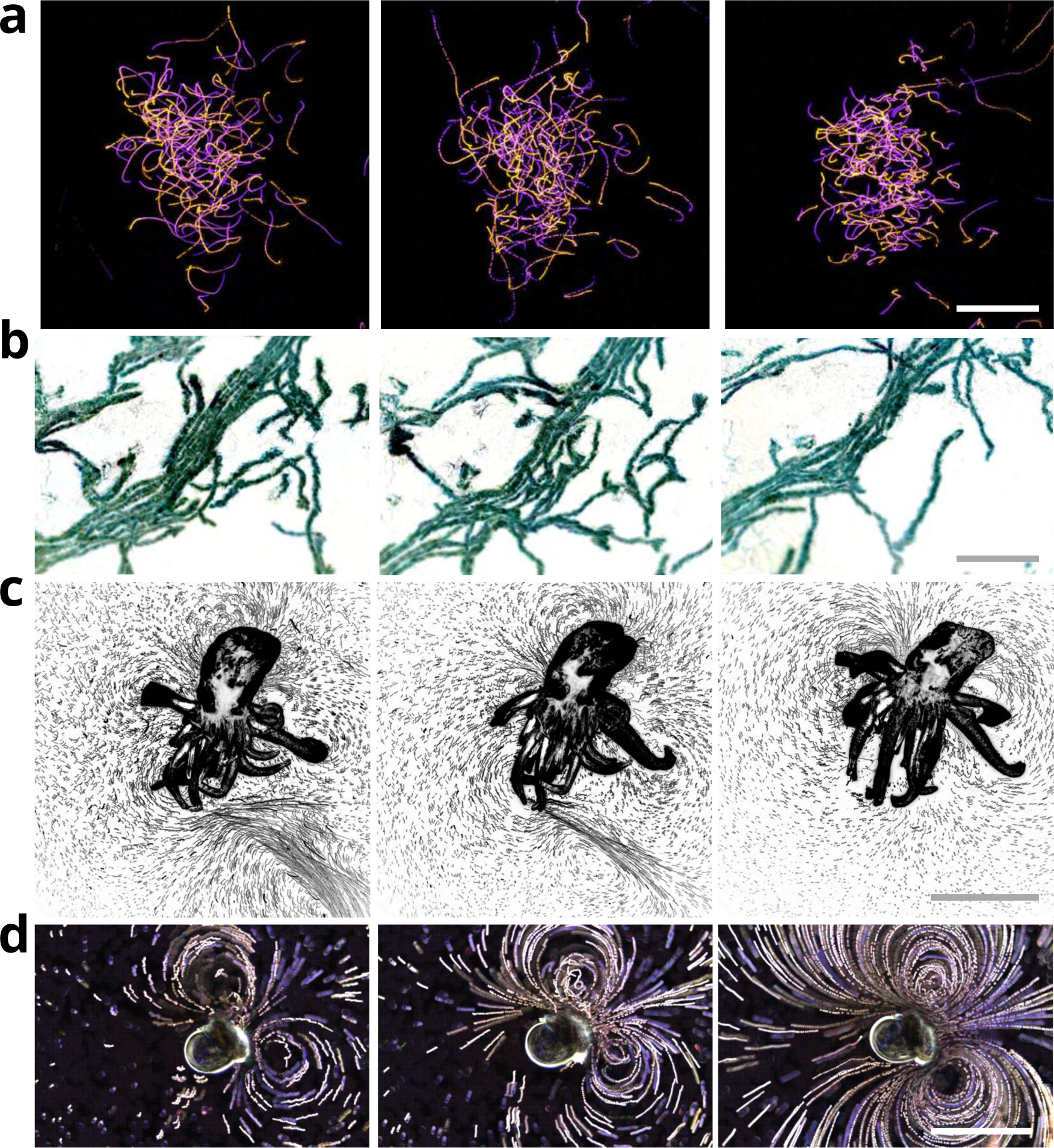
Application of Flowtrace to diverse datasets. (**a.**) Three frames from a movie of a flock of midges, with pathlines temporally color-coded from blue to orange (movie from ref. 6; *τ* = 333 ms, timepoints = 0, .66, 1.3 s, scale bar 60 mm).^28^ (**b.**) A crawling swarm of army ants (*τ* = 1 s, timepoints = 0, 1, 4.5 s, scale bar 20 mm). (**c.**) A sea anemone taking in a jet of water containing 6 *µ*m beads (*τ* = 4 min, timepoints = 0, 5.4, 18 min, scale bar 1 mm). (**d.**) A swimming moon snail larva, with 6 *µ*m beads mixed into the water (*τ* =2 s, timepoints = 0, 6, 20 s, scale bar 350 *µ*m).

## 5 Acknowledgements

The authors thank L. Y. Esherick and J. R. Pringle for providing anemones, and D. N. Clarke and C. J. Lowe for providing starfish larvae, moon snails, and microscopy equipment. This work was supported by a National Science Foundation Graduate Research Fellowship DGE-114747 (to W. G.), an ARO MURI Grant W911NF-15-1-0358 and an NSF CAREER Award (to M. P.).

## References

[1] Rohr, J., Hyman, M., Fallon, S. & Latz, M. I. Bioluminescence flow visualization in the ocean: an initial strategy based on laboratory experiments. Deep Sea Research Part I: Oceanographic Research Papers 49, 2009–2033 (2002).

[2] Bartol, I. K., Krueger, P. S., Jastrebsky, R. A., Williams, S. & Thompson, J. T. Volumetric flow imaging reveals the importance of vortex ring formation in squid swimming tail-first and arms-first. Journal of Experimental Biology 219, 392–403 (2016).

[3] Boot, M. J. et al. In vitro whole-organ imaging: 4d quantification of growing mouse limb buds. Nature Methods 5, 609–612 (2008).

[4] Lindken, R., Rossi, M., Große, S. & Westerweel, J. Micro-particle image velocimetry (µpiv): recent developments, applications, and guidelines. Lab on a Chip 9, 2551–2567 (2009).

[5] Shapiro, O. H. et al. Vortical ciliary flows actively enhance mass transport in reef corals. Proceedings of the National Academy of Sciences 111, 13391–13396 (2014).

[6] Han, S. J., Oak, Y., Groisman, A. & Danuser, G. Traction microscopy to identify force modulation in subresolution adhesions. Nature Methods 12, 653–656 (2015).

[7] Deforet, M. et al. Automated velocity mapping of migrating cell populations (AVeMap). Nature Methods 9, 1081–1083 (2012).

[8] Bharadvaj, B., Mabon, R. & Giddens, D. Steady flow in a model of the human carotid bifurcation. part iffflow visualization. Journal of biomechanics 15, 349–362 (1982).

[9] Santiago, J. G., Wereley, S. T., Meinhart, C. D., Beebe, D. & Adrian, R. J. A particle image velocimetry system for microfluidics. Experiments in fluids 25, 316–319 (1998).

[10] Eyal, S., Quake, S. R. et al. Velocity-independent microfluidic flow cytometry. Electrophoresis 23, 2653–2657 (2002).

[11] Merzkirch, W. Flow visualization (Elsevier, 2012).

[12] Miyake, R., Lammerink, T. S., Elwenspoek, M. & Fluitman, J. H. Micro mixer with fast diffusion. In Micro Electro Mechanical Systems, 1993, MEMS’93, Proceedings An Investigation of Micro Structures, Sensors, Actuators, Machines and Systems. IEEE., 248–253 (IEEE, 1993).

[13] Miles and, R. B. & Lempert, W. R. Quantitative flow visualization in unseeded flows. Annual review of fluid mechanics 29, 285–326 (1997).

[14] Bayraktar, T. & Pidugu, S. B. Characterization of liquid flows in microfluidic systems. International Journal of Heat and Mass Transfer 49, 815–824 (2006).

[15] Mercado, J. M. et al. Lagrangian statistics of light particles in turbulence. Physics of Fluids (1994-present) 24, 055106 (2012).

[16] Kertzscher, U., Berthe, A., Goubergrits, L. & Affeld, K. Particle image velocimetry of a flow at a vaulted wall. Proceedings of the Institution of Mechanical Engineers, Part H: Journal of Engineering in Medicine 222, 465–473 (2008).

[17] Stamhuis, E. & Videler, J. Quantitative flow analysis around aquatic animals using laser sheet particle image velocimetry. Journal of Experimental Biology 198, 283–294 (1995).

[18] Adrian, R. J. & Westerweel, J. Particle image velocimetry. 30 (Cambridge University Press, 2011).

[19] Stamhuis, E., Videler, J., van Duren, L. & Müller, U. Applying digital particle image velocimetry to animal-generated flows: Traps, hurdles and cures in mapping steady and unsteady flows in re regimes between 10–2 and 105. Experiments in Fluids 33, 801–813 (2002).

[20] Olsen, M. & Adrian, R. Out-of-focus effects on particle image visibility and correlation in microscopic particle image velocimetry. Experiments in fluids 29, S166–S174 (2000).

[21] Garcimart´in, A. et al. Flow and clogging of a sheep herd passing through a bottleneck. Physical Review E 91, 022808 (2015).

[22] Attanasi, A. et al. Information transfer and behavioural inertia in starling flocks. Nature physics 10, 691–696 (2014).

[23] Vicsek, T. & Zafeiris, A. Collective motion. Physics Reports 517, 71–140 (2012).

[24] Rau, K. R., Quinto-Su, P. A., Hellman, A. N. & Venugopalan, V. Pulsed laser microbeam-induced cell lysis: time-resolved imaging and analysis of hydrodynamic effects. Biophysical journal 91, 317–329 (2006).

[25] Hejnowicz, Z. & Kuczyńska, E. U. Occurrence of circular vessels above axillary buds in stems of woody plants. Acta Societatis Botanicorum Poloniae 56, 415–419 (1987).

[26] West, J. L. & Cameron, I. D. Using the medical image processing package, imagej, for astronomy. arXiv preprint astro-ph/0611686 (2006).

[27] Gilpin, W., Prakash, V. N. & Prakash, M. Vortex arrays and ciliary tangles underlie the feeding-swimming tradeoff in starfish larvae. arXiv preprint arXiv:1611.01173 (2016).

[28] Attanasi, A. et al. Collective Behaviour without Collective Order in Wild Swarms of Midges. PLoS Comput Biol 10, e1003697 (2014). URL http://dx.doi.org/10.1371%2Fjournal.pcbi.1003697.

[29] Gilpin, W., Prakash, V. N. & Prakash, M. Vortex arrays and ciliary tangles underlie the feeding-swimming tradeoff in starfish larvae. Bulletin of the American Physical Society 61 (2016).

[30] Shadden, S. C., Dabiri, J. O. & Marsden, J. E. Lagrangian analysis of fluid transport in empirical vortex ring flows. Physics of Fluids (1994-present) 18, 047105 (2006).

[31] Haller, G. Lagrangian coherent structures. Annual Review of Fluid Mechanics 47, 137–162 (2015).

